# Electrocorticographic responses to time-compressed speech vary across the cortical auditory hierarchy

**DOI:** 10.1101/354464

**Authors:** Ido Davidesco, Thomas Thesen, Christopher J Honey, Lucia Melloni, Werner Doyle, Orrin Devinsky, Oded Ghitza, Charles Schroeder, David Poeppel, Uri Hasson

**Author notes:** To whom correspondence should be addressed: Ido Davidesco, Psychology Department, New York University, New York, NY 10003.

## Abstract

Human listeners understand spoken language across a variety of rates, but when speech is presented three times or more faster than its usual rate, it becomes unintelligible. How the brain achieves such tolerance and why speech becomes unintelligible above certain rates is still unclear. We addressed these questions using electrocorticography (ECoG) recordings in 7 epileptic patients (two female). Patients rated the intelligibility of sentences presented at the original rate (100%), speeded rates (33% or 66% of the original sentence duration) and a slowed rate (150%). We then examined which parameters of the neural response covary with the transition from intelligible to unintelligible speech. Specifically, we asked whether neural responses: 1) track the acoustic envelope of the incoming speech; 2) “scale” with speech rate, i.e. whether neural responses elicited by slowed and speeded sentences can be linearly scaled to match the responses to the original sentence. Behaviorally, intelligibility was at ceiling for speech rates of 66% and above, but dropped significantly for the 33% rate. At the neural level, Superior Temporal Gyrus regions (STG) in close proximity to A1 (‘low-level’) tracked the acoustic envelope and linearly scaled with the input across all speech rates, irrespective of intelligibility. In contrast, secondary auditory areas in the STG as well as the inferior frontal gyrus and angular gyrus (‘high-level’) tracked the acoustic envelope and linearly scaled with input only for intelligible speech. These results help reconcile seemingly contradictory previous findings and provide better understanding of how information processing unfolds along the cortical auditory hierarchy.

## New & Noteworthy

The human brain can cope with large variations in speech rate. However, when speech is artificially accelerated, above a certain rate it becomes incomprehensible. This study investigated how the brain achieves this tolerance to speech rate, and what might constrain our understanding of speeded-up speech. Whereas in low-level auditory areas, neural responses scaled with speech rate irrespective of intelligibility, high-order brain regions could only track speech as long as it remained comprehensible.

## Introduction

Human listeners understand speech over a wide range of rates. Speech remains intelligible even when it is artificially slowed or accelerated up to 40% of its original duration (Dupoux and Green 1997; Mehler et al. 1993; Pallier et al. 1998; Sebastián-Gallés et al. 2000). However, how this tolerance to temporal variability is achieved at the neural level and why spoken language becomes unintelligible above certain rates is currently poorly understood.

Nourski et al. (2009) demonstrated that high-frequency (>70 Hz) electrocorticographic (ECoG) responses recorded directly from Heschl’s gyrus (A1) could track the speech envelope well outside of the intelligibility range. On the other hand, Ahissar et al. (2001) reported that time compression of speech beyond the intelligibility limit is associated with a sharp decrease in the temporal locking of auditory magnetoencephalographic (MEG) responses to the speech envelope. More recently, using functional MRI, Lerner et al. (2014) measured blood-oxygenation level dependent (BOLD) responses to speeded-up and slowed-down versions of a 7-minute narrative (50% to 200%). They found that for both the slowed-down and speeded-up rates, linearly scaled BOLD responses matched the response to the original narrative. This linear scaling of the neural responses was observed across the entire processing hierarchy, including early auditory regions as well as linguistic and extra-linguistic brain areas (but note that speech was always kept within the intelligibility range).

Although the findings described above seem to be contradictory, it is possible that they reflect different stages of processing along the auditory hierarchy. In a series of studies, we have demonstrated a neural hierarchy of Temporal Receptive Windows (TRWs) (Hasson et al. 2008; Honey et al. 2012; Lerner et al. 2011). Analogous to the notion of a spatial receptive field, TRW refers to the window of time in which information is being integrated. The TRW gradually increases from early sensory areas to higher-order perceptual and cognitive areas (Lerner et al. 2011). Therefore, we hypothesize that the short temporal integration windows of early auditory areas (e.g. A1) would enable the tracking of accelerated speech even outside of the intelligibility range. In contrast, in higher order areas, the integration of information may fail at high compression rates.

In the current study, we used ECoG recordings in seven neurosurgical patients to address the question of where along the cortical processing timescale hierarchy invariance to speech rate emerges. Participants were presented with a list of sentences spoken at a normal rate (100%) as well as slowed-down (150% duration) and speeded-up (66% and 33%) rates. Following Nourski et al. (2009) and Ahissar et al. (2001), we correlated the speech envelope with the envelope of the broadband (75-200Hz) neural responses at each speech rate. Based on the Lerner et al. (2014) study, we also tested the extent to which linear scaling of the neural responses elicited by speeded (or slowed down) sentences match the neural responses to the original speech rate. Whereas neural tracking is mostly sensitive to low-level properties of the speech signal (i.e. variations in amplitude across time), linear scaling can capture more high-level properties of speech processing (Lerner et al., 2014). Even though we did not have access to neural data from A1, we predicted that adjacent early auditory areas along the STG would exhibit speech rate invariance irrespective of intelligibility level. In contrast, areas outside of early auditory cortex, further along the processing hierarchy, which integrate sounds into intelligible syllables and words, would scale their neural activity with speech rate only within the intelligibility range.

## Materials and Methods

### Participants

Seven native speakers of English (2 female; 24-56 years old) experiencing pharmacologically refractory complex partial seizures were recruited via the Comprehensive Epilepsy Center of the New York University School of Medicine. Their clinical and demographic information is summarized in Table 1. Patients had elected to undergo intracranial monitoring for clinical purposes and provided written informed consent in accordance with New York University Medical Center Institutional Review Board. Electrode placement was determined based on clinical criteria without reference to this study. Patients had left-hemisphere (n=3), right-hemisphere (n=3) and bilateral (n=1) electrode coverage.

**Table 1:** Demographic and Recording Characteristics of Patients.

| Patient ID | Gender | Age (years) | WADA<sup>1</sup><br>language | Implanted<br>hemisphere | # of<br>implanted<br>electrodes |
| --- | --- | --- | --- | --- | --- |
| NY393 | M | 44 | Left | Left | 120 |
| NY339 | F | 24 | N/A | Left | 122 |
| NY415 | F | 56 | Left | Left | 124 |
| NY400 | M | 27 | Left | Bilateral | 124 |
| NY442 | M | 29 | N/A | Right | 64 |
| NY394 | M | 27 | Left | Right | 124 |
| NY451 | M | 25 | N/A | Right | 204 |

<sup>1</sup> Also known as intracarotid sodium amobarbital procedure; used to map language localization.

### Stimuli

A set of 33 spoken sentences with duration ranging between 3 and 3.3 seconds were selected from the Harvard sentences corpus (IEEE 1969). All sentences were recorded by a male speaker. Stimuli covered four speech rates: uncompressed (100%) speech, slowed (150%) and speeded speech (33% and 66% of the duration of the corresponding uncompressed signal; See Fig. 1A). Unfortunately, we could not include more intermediate speech rates because of the limited testing time available with each patient. The original rate and 33% conditions were represented by 33 sentences and the 66% and 150% conditions - by 25 sentences that were randomly selected from the set of 33. Sentences were presented consecutively, in pseudorandom order, until each sentence had been presented twice.

To control for sentence duration, we generated concatenated (C) sentences which were generated by (i) concatenating three different sentences and then (ii) time compressing the concatenated group by a factor of 3 (See Fig. 1A). Thus, each of these speeded sentence-groups had the same duration as one of the original sentences. The 8 sentences used to generate the 33%-concatenated (33C) condition were different than the ones used in the other conditions, and were sampled independently from the Harvard sentence corpus. Ten 33C sentences were generated in this manner, and each one of them was presented twice throughout the experiment, interleaved with the other conditions. Compression and dilation were performed using the Overlap-Add algorithm in Praat (Boersma and Weenink 2009), which preserves the spectral information of the uncompressed signal.

### Experimental design

Participants listened to a total of 252 sentences, divided into two blocks. Sentences were played at bedside by a laptop and speakers located in front of the patient. The experiment was controlled using Presentation^®^ software (Neurobehavioral Systems, Inc., Berkeley, CA, http://www.neurobs.com). Sentences were presented in a pseudo-random order, under the constraint that the same sentence was never repeated consecutively. The experiment was self-paced: following each sentences, patients verbally rated the intelligibility of the sentence they had just heard, using a 5 point scale from 1 (“not intelligible at all”) to 5 (“fully intelligible”).

### ECoG acquisition and preprocessing

Signals were recorded from 882 intracranially implanted subdural and depth electrodes (AdTech Medical Instrument Corp., WI, USA) in patients undergoing presurgical evaluation of pharmacologically intractable seizures. Electrode placement was determined solely on clinical grounds, and included grid (8×8 contacts), strip (1×4 to 1×12 contacts), and depth (1×8 contacts) electrode arrays with 10 mm inter-electrode spacing center-to-center (5 mm spacing in the depth electrodes). Neural signals were recorded on a Nicolet One EEG system, digitized at 512 Hz, and bandpass filtered between 0.5 - 250 Hz. Data were analyzed in MATLAB R2012a using custom scripts and the EEGLab toolbox (Delorme and Makeig 2004). At the preprocessing stage, each electrode was average-referenced by subtracting the mean voltage measured in all electrodes (Davidesco et al. 2013).

### Electrode localization

Magnetic Resonance (MR) anatomical images were obtained for each patient both before and after the implantation of electrodes. Electrodes were localized on the post-implant MR images using intraoperative photographs, manual identification, and a custom MATLAB tool based on the dimensions of the implanted electrode arrays (Yang et al. 2012). Next, the MR images were nonlinearly registered to MNI space using the DARTEL algorithm in SPM (Ashburner 2007), and the same transformation was used to map individual electrode coordinates into MNI space.

### Calculation of broadband power time courses

Broadband power fluctuations have been shown to reflect changes in population spiking activity (Crone et al. 2011; Manning et al. 2009; Nir et al. 2007; Whittingstall and Logothetis 2009). To compute broadband power time courses, Morlet wavelets (standard deviation 6 cycles) with center frequencies at 70, 75, 80, … 200 Hz were convolved with the voltage time series. Amplitude time series at line-noise frequencies of 120 and 180 Hz were discarded, leaving 25 distinct time series. Each individual amplitude time series was logarithmically transformed and then converted to a z-series by subtracting its mean and dividing by its standard deviation. The high frequency broadband power was then estimated as the mean of all 25 of the z-series and smoothed with a hamming window of 125ms (Honey et al. 2012).

### Statistical Analysis

For each one of the 882 electrodes and for each speech rate, two measures were computed:

#### 1) Neural tracking of the envelope of speech

Neural tracking was defined as the correlation between the speech envelope of a sentence and the corresponding broadband ECoG response.

The speech envelope was extracted for each sentence as follows: First, each sentence was filtered into sixteen critical bands logarithmically spaced between 230 and 3800 Hz; second, the Hilbert envelope was extracted for each band and then summed across bands. Finally, the resulting envelope time-course was down-sampled to 512 Hz to match it to the ECoG signal (Doelling et al. 2014). The ECoG broadband response was first shifted backwards by the response latency of each electrode (as estimated from the external localizer - see below). Then, to reduce any components that are not sentence-specific, the mean neural response across all sentences of a given speech rate was regressed out of each trial. To account for the difference in signal length across compression/dilation conditions, both the sentence envelope and the ECoG broadband responses were resampled to match the original sentence duration (i.e. 33% and 66% responses were up-sampled, 150% responses were down-sampled). Next, the first 300 ms and the last 300 ms were cropped from each trial in order to exclude onset or offset-related transients. Finally, the ECoG responses were averaged across the two repetitions of each sentence and correlated with the sentence envelope. Note that a correlation analysis was used, rather than a phase-locking analysis, because the latter requires multiple repetitions of each sentence (Luo and Poeppel 2007). Repeating the same time-compressed sentence multiple times can improve its intelligibility (Dupoux and Green 1997).

#### 2) Linear scaling

This analysis was used to test the extent to which linear scaling of the neural responses elicited by speeded or slowed down sentences match the neural responses to the original speech rate. In this analysis, the response to the original sentence (100%) was always used as a reference signal, and the neural response to each one of the other speech rates was resampled to match the original sentence duration (responses to speeded speech were upsampled, responses to slowed speech were down-sampled; See Fig. 3) (Lerner et al. 2014). Then, for every speech rate, the resampled neural response was correlated with the response to the original sentence. Note that the linear scaling analysis is expected to provide additional information, not captured by the speech tracking analysis. The speech envelope mainly reflects low-level properties of the speech signal (i.e. variations in amplitude over time). High-order cortical regions may no longer track the audio envelope of speech, but still be directly involved in speech processing (e.g. analyzing the grammatical structure of a sentence) (Honey et al. 2012). Therefore, linear scaling might be a more suitable measure to compare neural responses across low-and high-level cortical areas (see Discussion).

In all the analyses described above, the resulting correlation values were averaged across all sentences. A permutation test was used to assess the significance level of each electrode: sentence labels were randomly shuffled 1000 times, such that the neural response to a given sentence was correlated with the response to a different sentence. Then the empirical correlation value was compared to the null distribution of correlation values in order to assess the significance level of each electrode. FDR was used to correct for multiple comparisons (q<0.05).

### Selection of speech-specific electrodes

In the final analysis (Fig. 4), electrodes were selected based on a speech localizer task. This task allowed us to contrast the mean broadband power responses to speech and to noise. In the localizer task, patients viewed a still image depicting the lower part of a face, which was paired with a spoken word or a noise-vocoded word (Shannon et al. 1995). There were 20-30 trials of each type, and the patient was requested to press a button in response to a pre-defined target word. This task was part of another experiment on audiovisual speech. Due to the limited testing time available with each patient, this was the only dataset available for electrode selection.

However, the topography of speech-selective electrodes obtained based on this task was similar to that reported by previous studies that only used auditory stimuli (Edwards et al. 2009). For each trial the mean response in a time window of 50-500 ms following stimulus onset was computed. Then, a t-test was used to assess whether each electrode showed a significant difference in the response to speech and noise. In addition, a speech selectivity index was computed for each electrode as (see Fig. 4A):

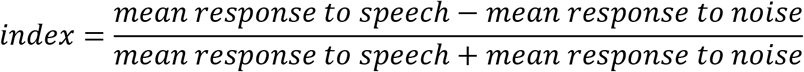

A total of 40 electrodes showed a significantly stronger response to speech compared to noise (False Discovery Rate (FDR) corrected, q<0.01), and were thus defined as “speech-specific” and used for subsequent analyses.

The localizer task also enabled us to extract the response latency of each electrode. The Student’s t-test was used to compare the broadband power response at each individual time point against a pre-stimulus baseline. The response latency was defined as the time, within the time series of broadband power, at which power first (i) became significantly larger than its prestimulus baseline value, and (ii) remained significantly higher than baseline for at least 10 successive sampling points (Davidesco et al. 2013). The averaged response latency of speech-selective electrodes was 150ms ± 54ms (mean ± standard deviation).

## Results

### Intelligibility

Patients rated the intelligibility of each sentence on a scale from 1 to 5 (where 5 is fully intelligible). Intelligibility was near ceiling for the 66%, 100% and 150% rates and, as expected, dropped sharply for the 33% and 33C conditions. Specifically, intelligibility significantly decreased from a level of 4.88±0.05 (mean rating ± standard error of the mean) for the original duration to a level of 2.78±0.38 for 33% (p=0.008; one-sided Wilcoxon’s signed-rank test), and to a level of 2.19±0.34 for 33C (p=0.008) (Fig. 1B).

**Figure 1:**
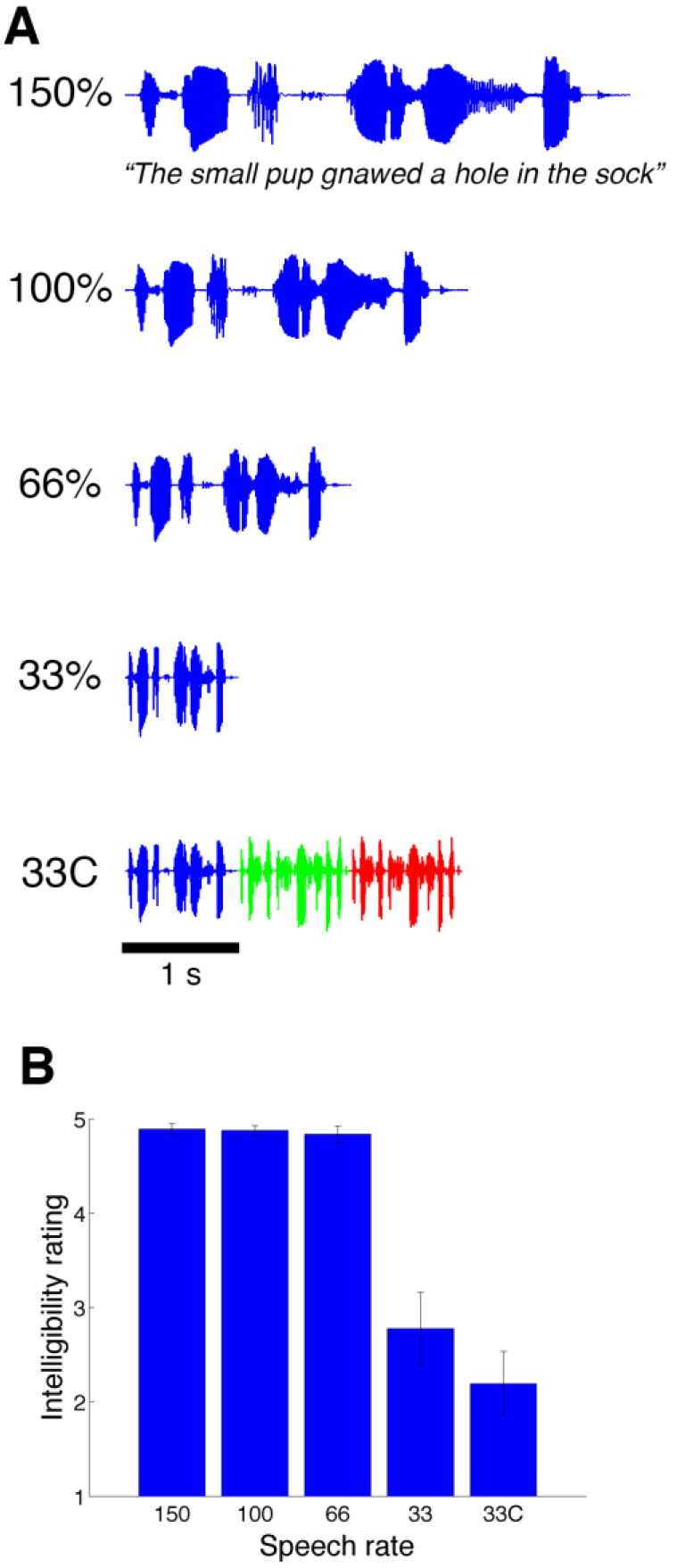
Experimental protocol and behavioral results. **A.** Experimental design: participants were presented with sentences at four different rates, with durations ranging from 33% to 150% of the original sentence duration. Participants also listened to sentences from the “33C” condition in which 3 different sentences were concatenated and then compressed by a factor of 3, to match the duration of the original sentence. **B.** Behavioral results (mean ± SEM): intelligibility ratings were at ceiling for rates of 66% and above and dropped dramatically for 33% compression.

### Neural tracking

We first assessed the extent to which neural responses tracked the audio envelope of each sentence. Figure 2A-B depict, for two different electrodes—low-level and high-level—the broadband responses (green) and audio envelope (red) for a single sentence at different speech rates. The low-level auditory electrode (Fig. 2A) closely tracked the audio envelope for all speech rates. In contrast, the high-level STG electrode (Fig. 2B) showed a sharp decrease in envelope tracking for the 33% and 33C conditions. To investigate the spatial topography of neural tracking of the speech envelope, we conducted a whole-brain analysis by assessing the significance of speech tracking across all recorded electrodes using a permutation test corrected for multiple comparisons (see Methods). Figure 2C shows the speech envelope tracking maps for each speech rate. Even though we did not have access to neural data from Heschl’s gyrus, seven electrodes, localized mainly along the lateral sulcus in the vicinity of early auditory areas, displayed significant speech tracking for the most compressed speech levels (33% and 33C). For intelligible speech rates (66% and above), as expected, most of the electrodes that showed significant speech tracking were clustered along the STG as well as in the inferior frontal gyrus. Slowed-down speech (150%) yielded the highest number of speech tracking sites, with some extending to the angular gyrus and supramarginal gyrus.

**Figure 2:**
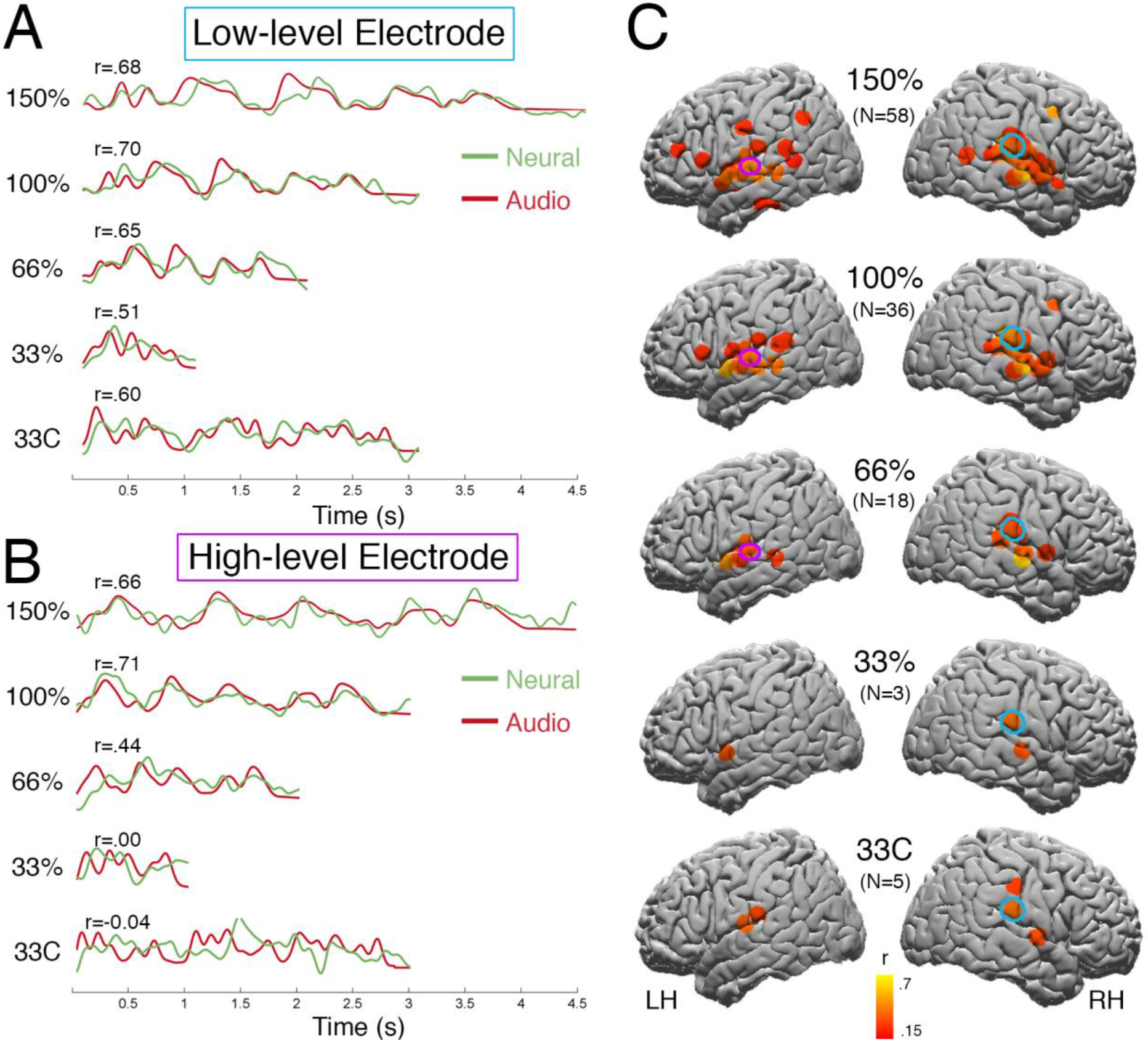
Speech envelope tracking. **A.** Broadband responses of an example low-level electrode (marked in cyan in panel C) to a single sentence presented at different rates (green) as well as the audio envelope of that sentence (red). Correlation values correspond to the single sentence that is depicted in panel A. **B.** Same as (A) for an example high-level electrode (marked in purple in panel C). **C.** Spatial distribution of all the electrodes that showed significant speech tracking (p < 0.05, FDR corrected) at each one of the speech rates. Electrodes are color-coded based on the correlation of the broadband response with speech audio envelope. Significance was assessed using a permutation test.

Could these results be driven by the difference in signal duration across conditions? To address that question, we compared the 33C and 100% conditions. These two conditions differ only by speech rate, not by signal length (see Methods). The 100% condition was characterized by widespread speech tracking along the anterior and posterior STG as well as the inferior frontal gyrus, whereas in the 33C condition, only 5 electrodes in the vicinity of early auditory areas exhibited significant speech tracking. Moreover, prior to measuring speech tracking, both the sentence envelope and the ECoG broadband responses for the compressed/dilated conditions were resampled to match the original sentence duration, therefore the number of time points in the neural response was held constant across speech rates (see Methods).

**Figure 3:**
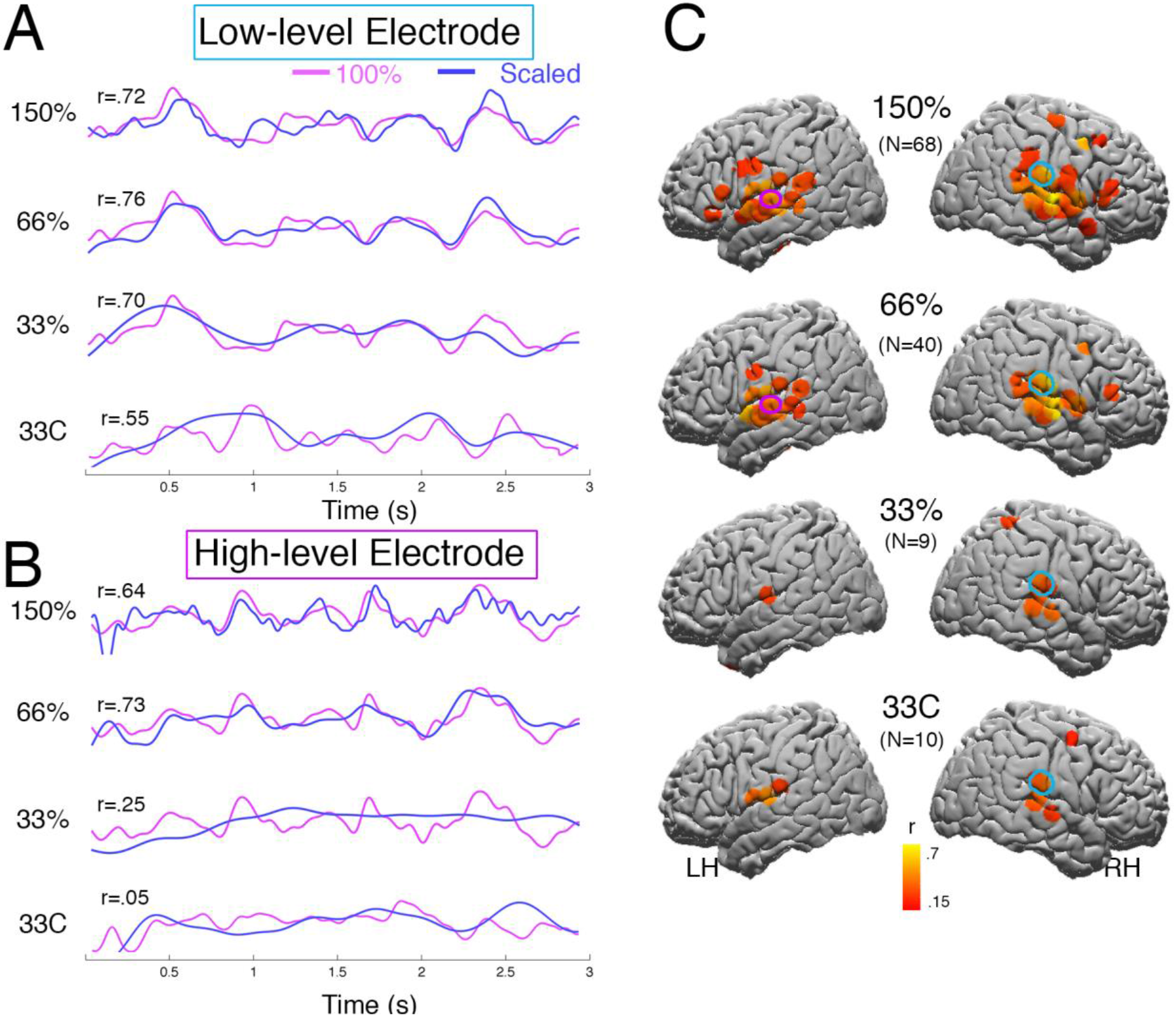
Temporal scaling. **A.** Broadband responses of an example low-level electrode (same electrode as in Fig. 2A). The neural responses to speeded (slowed) sentences (blue) were up-sampled (down-sampled) to match the length of the neural response to the original sentence (pink). **B.** Broadband responses of an example high-level electrode (same electrode as in Fig. 2B). **C.** Spatial distribution of electrodes showing significant temporal scaling (p<0.05; FDR corrected) for each speech rate.

### Linear scaling

Based on the work of Lerner et al. (2014), we also examined the linear scaling of the neural responses. Here, we measured the extent to which the responses for the speeded (slowed) sentences match the original (100%) by up-sampling (down-sampling) the neural responses (see Methods) (Lerner et al. 2014).

Figure 3A-B depict the scaled neural responses (blue) and the response to the original sentence that served as a reference (pink). For example, in the case of 150% (slowed) speech, the neural response was compressed (i.e. down-sampled) to match the 100% response, and then the two responses were correlated.

Similarly to the speech tracking analysis, we identified two representative electrodes. For low-level auditory electrodes (Fig. 3A), significant response scaling was observed across all speech rates, even outside the intelligibility range. In contrast, for high-order STG electrodes (Fig. 3B), temporal scaling was observed only for intelligible speech (66% and 150%) and not for unintelligible speech (33% and 33C). This step-like transition in temporal scaling from intelligible to non-intelligible speech was also evident in a whole-brain analysis. Significant temporal scaling along STG, Inferior Frontal Gyrus (IFG) and supramarginal gyrus was observed for intelligible speech (66% and 150%). In contrast, scaling of neural responses to unintelligible speech was mainly confined to STG sites in close proximity to early auditory cortex.

Finally, to further explore how speech rate affects neural processing in language related areas, we focused our analysis on 40 electrodes, which exhibited increased neural response to speech relative to non-speech stimuli, defined using an independent localizer task (see Methods). These electrodes were mainly clustered along the right and left STG, with the exception of 4 electrodes that were distributed over the IFG and motor cortex (Fig. 4A). Across the 40 speech-specific electrodes, the correlation between broadband responses and audio envelope (Fig. 4B) decreased monotonically as speech rate increased. Unlike speech envelope tracking, temporal scaling values dropped sharply, in a step-like function, for non-intelligible speech, in accordance with intelligibility ratings (compare Fig. 4C to Fig. 1B). To directly assess the transition from intelligible to non-intelligible speech across metrics, we conducted a two-way repeated measures ANOVA with speech ratio (66% vs. 33%) and metric (envelope tracking vs. temporal scaling) as factors. There was a highly significant interaction between these two factors (F(1,39)=46.80, p<10^−9^), indicating that temporal scaling dropped significantly in the transition between 66% and 33%, whereas speech tracking did not. This suggests that temporal scaling might be a more sensitive measure of speech processing that is more closely related to the observed behavioral effect.

**Figure 4:**
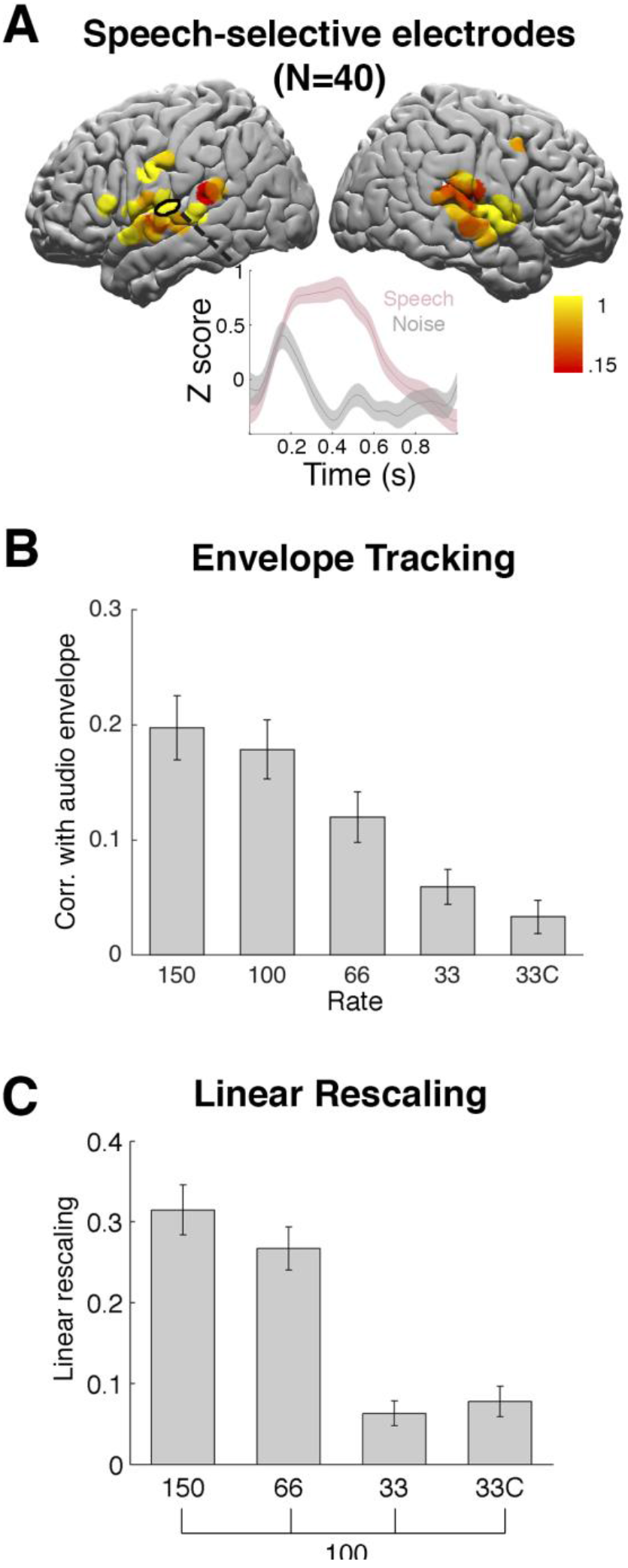
Response profile of speech-specific electrodes. **A.** Speech specificity map: electrodes that showed a significantly stronger response to individual words compared to noise-vocoded words in an independent localizer task (p < 0.05, FDR corrected). Electrodes are color coded according to speech specificity (0 = non selective, 1 = highly selective). Inset shows the broadband (70-200 Hz) responses to speech (red) and noise (black) of an example speech-selective electrode. **B.** Speech tracking correlation values (mean ± SEM) across 40 speech-specific electrodes as a function of speech rate. **C.** Same as (B) for linear scaling values.

## Discussion

The human auditory system can comprehend spoken language with remarkable tolerance to speech rate. Such tolerance, however, is limited. In particular, it has long been known that artificially time-compressing speech to a level beyond what is normally encountered in everyday listening (e.g., beyond compression by 3) hinders intelligibility at the word level and comprehension at the sentence level (Dupoux and Green 1997; Foulke and Sticht 1969; Garvey 1953; Ghitza 2014). Here, we recorded ECoG responses to sentences presented at speeded rates (33%, 66%), at the original rate (100%) and at a slowed rate (150% of the original sentence duration). Behaviorally, patients reported much lower understanding of highly compressed speech (33%; Fig. 1B). At the neural level, we observed two distinct response profiles: 1) A ‘low-level’ profile in electrodes along the STG, adjacent A1+, in which envelope tracking and linear scaling were observed across all speech rates; 2) A ‘high-level’ profile in electrodes further along the cortical hierarchy in anterior STG, posterior STG, IFG and Supramarginal gyrus, in which we observed significant speech tracking and linear scaling only within the intelligibility range (Figs. 2 and 3). Most of the electrodes in the current study exhibited the ‘high-level’ response profile, potentially as a result of the limited electrode coverage (i.e. we did not have access to recordings directly from Heschl’s Gyrus).

Our results help reconcile seemingly contradictory findings in the literature. Nourski et al. (2009), using intracranial recordings, demonstrated that Heschl’s gyrus (primary auditory cortex) can track the speech envelope well outside the intelligibility range. On the other hand, Ahissar et al. (2001), using MEG, reported that time compression of speech beyond the intelligibility limit is associated with a sharp decrease in speech envelope tracking. Our results suggest that these previous findings might correspond to distinct processing stages along the cortical processing hierarchy. Even though we did not have access to recordings in Heschl's gyrus within the Sylvian fissure, adjacent areas along the STG provided similar findings to these reported by Nourski et al. (2009). In contrast, the response in higher-order linguistic and extra-linguistic areas along the STG, IFG and supramarginal gyrus exhibited similar response profile to that reported by Ahissar et al. (2001) and Lerner et al. (2014). These results are consistent with a recent intracranial EEG study, which demonstrated a hierarchical organization of sound processing from the primary auditory cortex, where activity closely reflects the acoustic features of the stimulus, through the STG, where activity reflects both acoustic features and task demands, to the prefrontal cortex, which is mainly modulated by task requirements and behavioral performance (Nourski 2017). A related question that has received attention in the literature is where along the cortical hierarchy is the bottleneck in the processing of time-compressed speech. Our results, in accordance with Nourski et al. (2009), demonstrate that early auditory cortex can track speech outside of the intelligibility range, and it is therefore not to be considered the bottleneck. In accordance with our hypothesis, the first sites to track the speech envelope and to exhibit neural scaling only for intelligible speech were located along the STG. Interestingly, these areas seem to have an intermediate processing timescale in the order of few hundreds of milliseconds, which corresponds to the formation of syllables and the integration of syllables into words (Hasson et al. 2008; Honey et al. 2012; Lerner et al. 2011). Given that information flows upstream along the timescale hierarchy (from early auditory cortex to linguistic and extra-linguistic regions), it is reasonable to hypothesize that the bottleneck lies in areas with relatively short TRW that decode syllables and integrate syllables into words.

Once acoustic information is integrated into words, it can be transmitted up the timescale hierarchy to areas with longer TRW needed for the integration of words into sentences and sentences into paragraphs. Indeed, we observed that the neural activity in high-level linguistic areas in the IFG and supramarginal gyrus tracked the speech envelope only for intelligible speech. These findings are congruent with an fMRI study that demonstrated that the inferior frontal gyrus and the superior temporal sulcus show an invariant response to moderate compression rates followed by a sharp decline in activation for non-intelligible compressed speech (Vagharchakian et al., 2012). The current study extends these findings by demonstrating that millisecond-by-millisecond STG responses track the speech envelope and linearly scale with speech rate, as long as speech remains intelligible. Furthermore, our finding that language areas scale their dynamics in response to speech rate suggests that temporal integration windows should also be assessed using relative information-based units (e.g. the number of syllables) rather than merely in absolute temporal units (e.g. milliseconds), which vary across compression rates (Lerner et al. 2014).

It is worth noting that due to the limited testing time available with neurosurgical patients, this study only examined four speech rates. In future studies, it would be informative to sample the intelligibility spectrum more densely in order to map the transition from intelligible to non-intelligible speech.

Why do areas with an intermediate TRW, which are presumably involved in syllable formation and the integration of syllables into words, fail to track speech compressed by a factor of 3 or more? A potential explanation is provided by TEMPO (Ghitza 2011), a model that epitomizes recently proposed oscillation-based models of speech perception (Ahissar and Ahissar 2005; Ding and Simon 2009; Ghitza and Greenberg 2009; Giraud and Poeppel 2012; Hyafil et al. 2015; Lakatos et al. 2005; Peelle and Davis 2012; Poeppel 2003). TEMPO postulates a cortical computation principle by which decoding is performed within a hierarchical time-varying window structure, synchronized with the input on multiple time scales. The windows are generated by a segmentation process, implemented by a cascade of oscillators, governed by the theta oscillator, which provides syllabic segmentation. These oscillators operate within a constrained range of frequencies (the biophysical frequency range of theta). Critically, intelligibility remains high as long as theta is in sync with the input (as is the case for moderate speech speeds) and it sharply deteriorates once theta is out of sync (when the input syllabic rate is outside the theta frequency range). The notion that cortical oscillations are closely related to speech uptake capacity has received support from several recent studies (Borges et al. 2018; Pefkou et al. 2017). The findings of the current study suggest that the neuronal circuitry of the theta oscillator might be located at the STG level.

Finally, it is worth noting the difference between the insights provided by the neural tracking and the linear scaling measures. While neural tracking measures how well the neural response matches the acoustic envelope, linear scaling captures how consistent the neural response is across speech rates. Neural tracking is mostly sensitive to low-level properties of the speech signal (i.e. variations in amplitude across time). Whereas low-level regions (e.g. A1+) are expected to closely track the speech envelope, this might not be the case with high-order cortical regions. Indeed, Honey et al. (2012) demonstrated that regions with long TRW (e.g. medial frontal gyrus) no longer track the audio envelope, and yet respond very reliably to audiovisual stimuli. In the current study, more electrodes showed significant linear scaling compared to speech tracking (e.g. in the 66% rate: 18 electrodes compared 40 electrodes, respectively). Moreover, note that the degree of linear scaling dropped sharply with speech rate, in a better correspondence to the behavioral data, compared to a shallower drop in neural tracking (Fig. 4). This finding extends Lerner et al. (2014), who observed a linear scaling effect in temporally sluggish fMRI measurements. Here, we show that millisecond-by-millisecond neural responses recorded directly from high-level auditory regions can linearly scale with speech rate, as long as speech remains intelligible. Our findings suggest that neural tracking at secondary auditory areas in the STG, and beyond, is a prerequisite for intelligibility. As long as envelope tracking is maintained, the linearly scaled neural responses are remarkably stable as speech rate varies, and speech is intelligible.

## Acknowledgements

This work was supported by US National Institutes of Health grant R01 MH094480 (U.H., C.J.H.). We would like to acknowledge the contribution of the patients who participated in this study.

## Disclosures

The authors declare no competing financial interests.

